# Strengths of relationships among soil microbial and organic matter properties are scale-dependent

**DOI:** 10.1101/2025.07.08.663705

**Authors:** Eva Simon, Ksenia Guseva, Lauren Alteio, Christina Kaiser

## Abstract

Relationships among variables in ecological systems are inherently scale-dependent, especially in heterogeneous systems. Yet it remains to be examined whether relationships among variables vary across observation scales in soil. Generally, it is desirable that observation scale matches the intrinsic scale of a process or pattern. Millimetre-sized soil aggregates are closer to the intrinsic scale of microbial communities than traditionally studied bulk soil samples, making them more suitable for studying potential links between microbial communities and their environment.

To explore the effect of observation scale on relationships among soil parameters, we measured bacterial, archaeal, and fungal taxa richness and density, organic matter properties (e.g., carbon and nitrogen content, stoichiometric and isotopic ratios), and soil water content in individual aggregates and aliquots of homogenised soil cores, bulk soil samples, in two soil layers. We analysed pairwise correlations among these variables and assessed whether individual aggregates systematically differed from bulk soil samples.

Organic matter properties were more strongly correlated in bulk soil samples, consistent with the idea that increasing the sample volume reduces noise. In contrast, microbial community and organic matter properties showed weaker correlations in bulk soil samples than aggregates in topsoil. In addition, we found that aggregates and bulk soil samples differed systematically in individual microbial and organic matter properties, particularly in the topsoil.

Our study demonstrates that relationships among variables in soil are spatial scale-dependent. Aggregates offer valuable insights into microbial communities in soil, complementing bulk soil samples, and are useful for studying links between microbial communities and their environment.

## 1 Introduction

Soil bacteria, archaea, and fungi drive large-scale biogeochemical cycles (Fierer, 2017). They mediate soil fertility, guarantee food security and manage the largest terrestrial carbon storage (Van Der Heijden et al., 2008; Fierer, 2017; Liang et al., 2017; Wilpiszeski et al., 2019). Disentangling which factors predominantly shape structure and activity of microbial communities and their biogeochemical process rates in soils is an ultimate, yet, still not fully achieved goal. To truly understand a process or pattern, “intrinsic scale”, the scale at which pattern or process operates, and “observation scale”, the scale of measurement or sampling, need to match (Wiens, 1989; Wu and Li, 2006; Wu et al., 2006). A mismatch of intrinsic scale and observation scale may even lead to erroneous conclusions (Wu and Li, 2006; Wu et al., 2006) and result in “greater confusion than enlightenment” (Wiens, 1989). If we want to understand how microbial communities and properties of their immediate environment are related, samples must meet or at least approximate the intrinsic scale at which microbes interact with their environment.

In all ecological systems, relationships among variables are assumed to be spatial scale-dependent (Wiens, 1989; Wu and Li, 2006). The spatial scale-dependency of phenomena like patterns and processes increases the more heterogeneous a system is (Wu, 2007). Soil is highly heterogeneous across several scales (Or et al., 2007; Young et al., 2008; Vos et al., 2013; Tecon and Or, 2017). Pore sizes, gases, liquids, and organic matter are heterogeneously distributed throughout the soil matrix, even at the micrometre- to millimetre-scale. This heterogeneity generates a vast number of microhabitats which provide numerous ecological niches to microbes (Or et al., 2007; Young et al., 2008; Vos et al., 2013; Tecon and Or, 2017). Bacterial cells have been observed to distribute heterogeneously throughout the soil pore space and cluster in micro-scale patches (Nunan et al., 2002, 2003). Overall, soil has been found to be sparsely populated. Estimations reach from considerably less than 1% of the soil surface being colonised by microorganisms (Young and Crawford, 2004) up to 1 to 5.8% of the available habitable surface area being colonised by microorganisms (Schmidt et al., 2024).

Soil contains aggregates, that are micro- to millimetre-sized clumps of soil whose particles cohere more strongly among themselves than with neighbouring particles (Martin et al., 1955). Soil aggregates are thought to form naturally in-situ, often when organic matter stimulates microbial activity. They are stabilised by various microbial mechanisms such as the microbial release of binding agents (like extracellular polysaccharides), physical enmeshment by fungal hyphae or actinobacteria, and increased aggregate hydrophobicity by microbial activity (Chenu and Cosentino, 2011). When binding forces become weaker over time, aggregates eventually disintegrate in response to external stresses such as drying-rewetting (Tisdall and Oades, 1982; De Gryze et al., 2005; Wang et al., 2019; Yudina and Kuzyakov, 2023; Garland et al., 2024). While not much is known about aggregate turnover times in a broad variety of soils and conditions, the average time for aggregate stability has been estimated to be between 4 and 95 days in selected soils (Plante et al., 2002; De Gryze et al., 2005).

We propose that individual soil aggregates are useful study objects for linking microbial community parameters and soil organic matter properties. Analysing such small-volume soil units from undisturbed soils individually may provide new insights as they approximate a spatial scale relevant to micrometre-sized microorganisms (Bailey et al., 2012, 2013; Wilpiszeski et al., 2019; Alteio et al., 2021; Simon et al., 2024). Analysis of individual millimetre-sized aggregates has already revealed that the composition of microbial communities varies strongly at the millimetre-scale, even across aggregates which are only a few centimetres-apart (Szoboszlay and Tebbe, 2021; Simon et al., 2024). Moreover, community properties like bacterial and fungal taxon richness have been found to be linked with organic matter properties such as carbon content at the aggregate-scale (Simon et al., 2024). The value of using individual aggregates as study units for linking microbial and organic matter properties is supported by the argument that soil aggregates offer specific micro-environments that differ in physical, chemical, and biological properties from the surrounding soil matrix (Rillig et al., 2017; Wang et al., 2019) and that they may represent microbial habitats that are stable for long enough to allow the establishment of relationships between microbial communities and their immediate surroundings (Rillig et al., 2017; Simon et al., 2024). This view of aggregates, however, has been recently challenged by the proposal that aggregates are not intrinsically formed soil units, but instead are arbitrary soil fragments with no qualitative difference to the surrounding soil matrix (Kravchenko et al., 2019; Vogel et al., 2022).

Microbial communities are traditionally investigated in homogenised soil samples with the aim to obtain samples that are representative for a bigger soil volume or sampling area. These samples are aliquots of soil samples which are produced by sieving and mixing one or several soil core(s), each comprising several cubic centimetres of soil. However, through this homogenisation all information about the small-scale spatial heterogeneity of the soil matrix is lost and a vast number of physico-chemically variable microhabitats and associated microbial communities, which were spatially separated from each other in the intact soil matrix are blended (Nunan, 2017; Smercina et al., 2021).

The differences between homogenised larger-volume soil samples and individual aggregates are that they (i) represent different volumes of soil, and that (ii) the sum of all aggregates may not be 100% representative of larger-volume soil samples if they also contain a non-aggregate fraction which is qualitatively different. The aim of this study was to determine whether potential relationships between microbial communities and organic matter properties can be better identified at the level of individual millimetre-sized aggregates or homogenised larger-volume soil samples. On the one hand, we expect that relationships between microbes and organic matter arise at microbial-relevant scales, which are better approximated by millimetre-sized soil samples. On the other hand, it is known from scale theory that larger-volume samples can potentially exhibit stronger correlations among variables than smaller-volume samples, even if the origin of the relationship is at the small scale. This is because larger-volume samples average out local heterogeneity, which has been proposed to reduce noise and potentially strengthen correlations among variables (Wiens, 1989). However, whether relationships between microbial and abiotic parameters are really scale-dependent in soil has yet to be tested.

In this study, we addressed two main questions: (i) Are individually collected aggregates representative of larger soil samples in terms of their microbial community, organic matter properties, and soil water content? Or do they represent a fraction with specific properties? (ii) Do individually collected 2-millimetre-sized aggregates and homogenised larger-volume soil samples depict the same relationships among and across microbial community, organic matter properties, and soil water content? Or are these relationships scale-dependent? To answer these questions, we (i) examined whether individual aggregates taken from larger-volume soil cores systematically differed in microbial parameters, soil organic matter properties, and soil water content from the homogenised rest of the soil cores (termed “composite soil samples” from now on) and (ii) compared correlations among microbial community, organic matter properties, and soil water content across individual soil aggregates and composite soil samples. For this, we concomitantly measured carbon and total nitrogen content, carbon-to-nitrogen ratio, isotopic ratio of carbon and nitrogen, soil water content, bacterial and archaeal, and fungal taxa richness and gene copy numbers from the same samples, which were either individual aggregates or composite soil samples. These samples were collected from a Beech forest soil in two soil depths. We focused on organic matter properties as organic matter and microbial communities are proposed to be closely linked.

## 2 Material and Methods

### 2.1 Study site and soil sampling

We sampled soil cores across a 11 x 65 metre-large rectangular area in a *Fagus sylvatica*-dominated temperate forest in Austria (48.11925 N, 16.04343 E, 580 m altitude), in Autumn 2020. 20 soil cores (3-centimetre diameter, 5-centimetre height) were taken in 0-5 centimetres soil depth and 20 more in 15-20 centimetres underneath topsoil cores. For more details on soil sampling see Simon et al (Simon et al., 2024). We hand-picked four to five individual approximately 2-millimetre-sized aggregates from each soil core and sieved remaining field-moist soil through a 2-millimetre-mesh, producing a “composite soil sample” for each soil core. With “aggregate” we refer to millimetre-sized clumps of soil that did not break apart but remained intact during the process of extracting soil from cores, transferring it to a petri dish, and hand-picking aggregates with tweezers, indicating that bonds within the soil clump were stronger than bonds with adjacent particles. Shortly, to collect individual soil aggregates, we transferred less than a teaspoon of soil from soil cores to a petri dish and picked soil aggregates which were approximately two millimetre in diameter with a fine tweezer. The petri dish contained a moist filter paper to avoid drying of soil aggregates during their handling and was closed between picking individual aggregates. We weighed intact aggregates individually on a fine balance directly after selecting them. Subsequently, we transferred them to individual safe-lock Eppendorf tubes, stored them at −80°C, freeze-dried them for 48 hours, weighed them again to determine their dry weight, and homogenised them by hand with a ball tool (which usually is used as a cake decorating tool) to obtain a homogenised sample. Individual aggregates and composite soil samples are representative for markedly different soil volumes (Fig. S1, composite soil sample: V=35 cm^3^, aggregate: V=0.0042 cm^3^).

### 2.2 Bacterial and archaeal, and fungal community variables

DNA was extracted from aliquots of homogenised, freeze-dried aggregates (0-5 cm: n=92, 3.52 ± 1.7 mg dry soil, 15-20 cm: n=98, 4.1 ± 1.9 mg dry soil) and freeze-dried composite soil samples (0-5 cm: n=20, 186 ± 15.3 mg dry soil, 15-20 cm: n=20, 209 ± 45.1 mg dry soil) with the Qiagen PowerSoil Pro Kit (Qiagen, Valencia, CA, USA) following the manufactureŕs protocol, except that we replaced the 10-minute vortexing step with a bead-beating step (30 seconds) on a FastPrep™-24 Classic Bead Beater (MP Biomedicals, Germany) for improved cell lysis.

We carried out a two-step PCR barcoding approach to amplify the V4 region of the 16S rRNA gene targeting bacteria and archaea and the fungal internal transcribed spacer region 2 (ITS2) of the nuclear rRNA operon(Pjevac et al., 2021). We used primers 515F (Parada et al., 2016) and 806R (Apprill et al., 2015) for the bacterial and archaeal 16S rRNA gene, and primers ITSOF-T (Taylor and McCormick, 2008; Tedersoo et al., 2008) and ITS4 (White, T. J. et al., 1990), and gITS7 (Ihrmark et al., 2012) and ITS4 (White, T. J. et al., 1990) in a consecutive PCR for fungal ITS2 region. Please see detailed PCR cycling conditions in Table S1. Sequencing of amplicons was performed on an Illumina MiSeq platform (V3 chemistry, 600 cycles) at the Joint Microbiome Facility of the Medical University of Vienna and the University of Vienna under the project ID JMF-2011-I. Raw sequencing data underwent amplicon pool extraction using the FASTQ workflow in BaseSpace (Illumina, default settings). PhiX contamination was removed using BBDuk (BBtools)(Bushnell, 2015), and demultiplexing of data was carried out using the demultiplex package (Laros JFJ, github.com/jfjlaros/demultiplex), allowing for one mismatch for barcodes and two for linkers and primers. Amplicon sequence variants (ASVs) were then inferred utilizing the DADA2 package (v. 1.18.0 (Callahan et al., 2016)) with default settings. Classification of 16S rRNA gene ASVs was performed against the SILVA database (SSU RefNR 99, release 138.1) using the classifier integrated into DADA2. Fungal ITS2 ASVs were classified against the UNITE database (v. 04.02.2020 (Abarenkov et al., 2020)).

#### 2.2.1 Bacterial and archaeal, and fungal richness

We determined bacterial and archaeal, and fungal richness (number of unique ASVs per sample) of individual aggregates and composite soil samples from rarefied datasets. For this, we randomly rarefied samples to 8185 for bacteria and archaea and 8195 sequences for fungi, only considering samples with at least 8000 sequences for calculation of ASV richness.

#### 2.2.2 Bacterial and archaeal, and fungal density

We measured DNA content of DNA extracts with a HS dsDNA assay following the manufacturer’s protocol on the Qubit 4.0 fluorometer (Thermo Fisher Inc, Waltham, MA, United States). We determined 16S and ITS1 gene copy numbers in DNA extracts via the QX200 Droplet Digital PCR (Bio-Rad). See Supplement for primers and PCR cycling conditions (Table S2). We diluted DNA extracts (if necessary) to 0.1 ng (16S rRNA genes) or 0.5 ng (ITS1 region) of total DNA per PCR reaction to optimize the separation of negative and positive droplets. We analysed droplet readings (>10000 droplets, ≥250 positive droplets, ≥250 negative droplets) using the QuantaSoft software (Bio-Rad). Finally, we calculated gene copy number per gram dry soil as a proxy of microbial density.

### 2.3 Abiotic variables

#### 2.3.1 Organic matter properties: Organic carbon and nitrogen content, stoichiometric and isotopic ratios of carbon and nitrogen

We measured organic carbon and nitrogen content, isotopic ratios of carbon (δ^13^C) and nitrogen (δ^15^N) of aliquots of homogenised, freeze-dried aggregates (0-5 cm: n=92, 0.5 ± 0.04 mg dry soil, 15-20 cm: n=98, 0.6 ± 0.03 mg dry soil) and of milled, freeze-dried composite soil samples (0-5 cm: n=20, 8.29 ± 0.39 mg dry soil, 15-20 cm: n=20, 12.1 ± 0.47 mg dry soil) via an elemental analyser (EA IsoLink CN, Thermo Scientific) coupled to an isotope ratio mass spectrometer (Delta V Advantage Isotope Ratio MS Hi Pan CNOS, Thermo Scientific) via a ConFlo IV interface (Thermo Scientific). We calculated the stoichiometry ratio of carbon to nitrogen (carbon-to-nitrogen) and report stable isotope compositions in delta notation (δ^13^C, δ^15^N) in parts per thousand (‰) relative to the international standard VPDB, where d=1000/[(R_sample_/R_standard_)-1], where R represents the ratio of ^13^C/^12^C and ^15^N/^14^N, respectively.

#### 2.3.2 Soil water content

We determined gravimetric soil water content of individual aggregates (0-5 cm: n=64, 15-20 cm: n=73) and composite soil samples (0-5 cm: n=20, 15-20 cm: n=20) by subtracting weight of field-moist soil from weight of freeze-dried (48 h) intact aggregates and composite soil samples.

The number of measurements differs among sample types and variables (Table S3).

### 2.4 Statistical analysis

All data analyses were performed in R (v 4.2.1) (R Core Team, 2022). We used the package “TreeSummarizedExperiment” (v 2.4.0) (Huang et al., 2021) for microbiome data handling.

We utilised principal component analysis (PCA) based on the carbon and nitrogen content, δ^13^C and δ^15^N, and carbon-to-nitrogen ratio of samples (aggregates: n=190, composite soil samples: n=40) to identify potential clustering of samples by soil depth and/or sample type. Before performing PCA, all variables were centred and scaled to achieve standardized and comparable variables.

We performed Kruskal Wallis tests to detect significant differences (p<0.05) in carbon and nitrogen content, carbon-to-nitrogen ratio, δ^13^C and δ^15^N, and soil water content between aggregates (n=190) and composite soil samples (n=40) of the same soil depth, respectively.

One of the aims of this study was to test whether sample volume (i.e. grain size) affects the observed strength of the relationship between two given parameters. We examined the strengths of the relationships between C, nitrogen content, carbon-to-nitrogen ratio, isotopic ratios (δ^13^C, δ^15^N), bacterial and archaeal richness, fungal richness, and microbial densities in aggregates and composite soil samples by examining pairwise Spearman correlation coefficients as well as the significance of the correlations (P value). For composite soil samples, we computed Spearman correlation coefficients and P values for each parameter combination in each soil depth (n=20). To keep the sample extent (20 samples per soil layer) constant and only assess differences caused by a change in sample grain (volume), we calculated correlation coefficients and significance of correlations across the same number of aggregates (20 samples per soil layer). To cover a range of possible aggregate combinations in our dataset, we created 5000 combinations of 20 aggregates per soil depth by randomly selecting one aggregate from each soil core. We then computed pairwise Spearman correlation coefficients for each subset of possible combinations and calculated the median from all correlation coefficients of the same soil depth. We also calculated the median of all P values of the obtained 5000 correlations for each parameter combination.

As small-volume samples have a higher inter-sample heterogeneity compared to larger-volume samples it may be useful to increase sample numbers correspondingly. In order to assess whether the identification of potential links between communities and organic matter properties can be improved by including a higher number of small-scale samples we determined correlations across all sampled individual aggregates (i.e. 90-100 per soil depth), and compared them with all composite soil samples (i.e. 20 per soil depth) (Fig. S2). We visualised correlation coefficients in correlation heatmaps using the “corrplot” package (v 0.92) (Taiyun Wei and Viliam Simko, 2017). To illustrate whether correlations were stronger in aggregates or composite soil samples we subtracted absolute values of correlation coefficients of composite soil samples from (median) coefficients of aggregates and visualised them as heatmaps.

## 3 Results

### 3.1 Individual soil aggregates were systematically different from composite soil samples

#### 3.1.1 Aggregates differed in their chemical composition from composite soil samples

We performed principal component analysis (PCA) based on carbon (C), nitrogen (N) content, δ^13^C, δ^15^N, and carbon-to-nitrogen ratio of samples to assess (dis)similarities in chemical composition between aggregates and composite soil samples in the two soil depths. Within each sample type, chemical composition was influenced by soil depth (mostly along PC1, which explained 80.3% of the total variability). At a given soil depth, aggregates and composite soil samples formed partially overlapping but still distinguishable clusters, with the aggregate clusters being consistently shifted towards higher δ^13^C and lower carbon-to-nitrogen ratios compared to the composite soil clusters. Topsoil aggregates were most diverse in their chemical composition, overlapping with composite soil samples from both soil layers (Fig. 1).

**Fig. 1.**
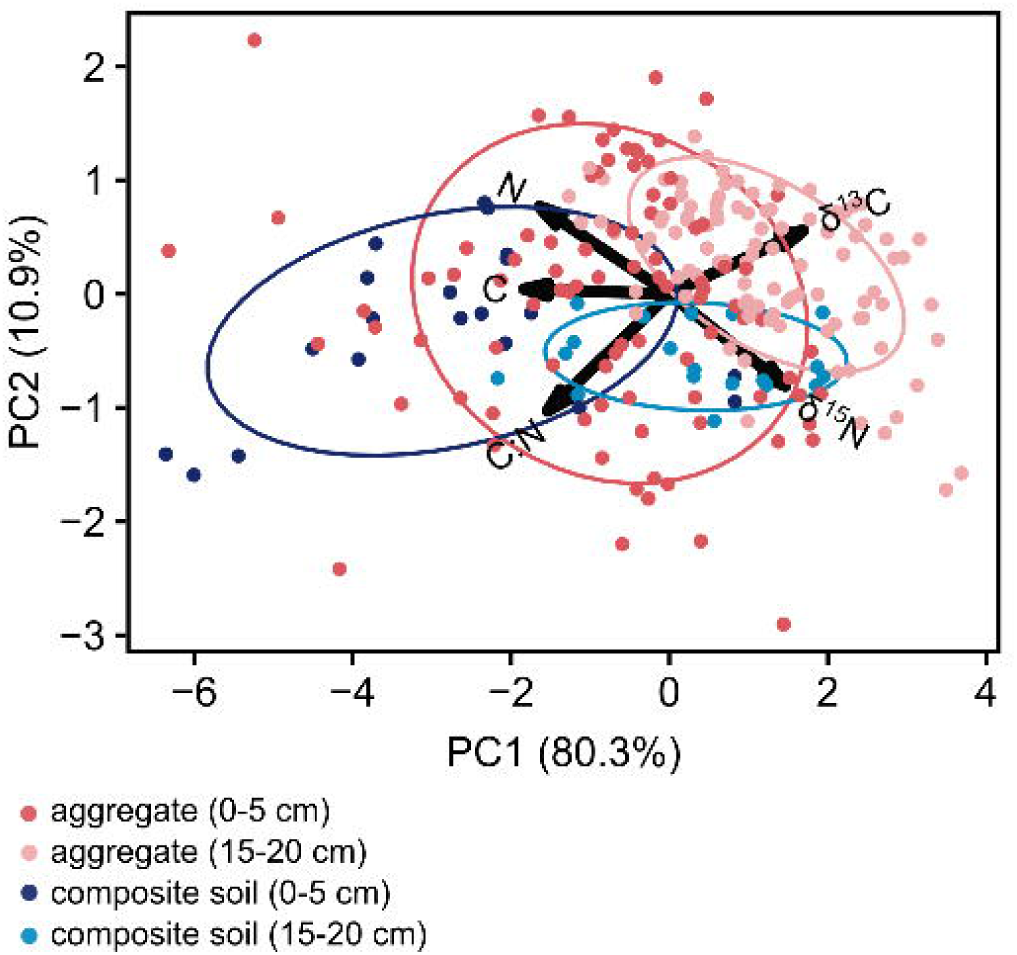
Aggregates and composite soil samples differed systematically in their chemical composition. PCA plots show the clustering of aggregates and composite soil samples in 0-5 and 15-20 centimetres soil depth based on their carbon (C) and nitrogen (N) content, δ^13^C, δ^15^N, and carbon-to-nitrogen ratio (C:N) (aggregates: 0-5 cm: n=92, 15-20 cm: n=98, composite soil: 0-5 cm: n=20, 15-20 cm: n=20). The first two principal components (PCs), which explain most of the variation, are plotted. Each point represents a sample and each vector (arrow) corresponds to a variable. Arrows point in the directions of the greatest increase in each variable. Variables that are perfectly correlated to an axis are represented by vectors that are parallel to that axis. The angle between vectors shows the strength and direction of the correlation between these variables: a small angle indicates strong positive correlation, whereas vectors pointing in opposite directions indicate strong negative correlation of variables. Ellipses encompass 68% of the samples.

#### 3.1.2 Aggregates had lower carbon and nitrogen content than composite soil samples in topsoil, were ^13^C-enriched, and exhibited lower carbon-to-nitrogen ratio than composite soil samples in both soil layers

We compared abiotic and microbial parameters between individual 2-millimetre-sized aggregates (n=190) and composite soil samples (n=40) for 0-5- and 15-20-centimetres soil depth to assess systematic differences between the two sample volumes. Carbon (Fig. 2A) and nitrogen (Fig. 2B) content were significantly lower in aggregates than in composite soil samples in topsoil (C: χ^2^=16.4, P value≤0.0001, N: χ^2^=13.1, P value≤0.0003), but was similar in 15-20 centimetres soil depth. In contrast, aggregates were more enriched in ^13^C compared to composite soil samples (Fig. 2D, 0-5 cm: χ^2^=30.2, P value≤0.0001, 15-20 cm: χ^2^=30.1, P value≤0.0001) and were characterised by a lower carbon-to-nitrogen ratio than composite soil samples in both soil depths (Fig. 2C, 0-5 cm: χ^2^=8.9, P value=0.003, 15-20 cm: χ^2^=22.2, P value≤0.0001). Topsoil aggregates were also enriched in ^15^N compared to composite soil (Fig. 2F, χ^2^=8.2, P value=0.004). Soil water content was similar for aggregates and composite soil samples in both soil depths (Fig. 2F).

**Fig. 2.**
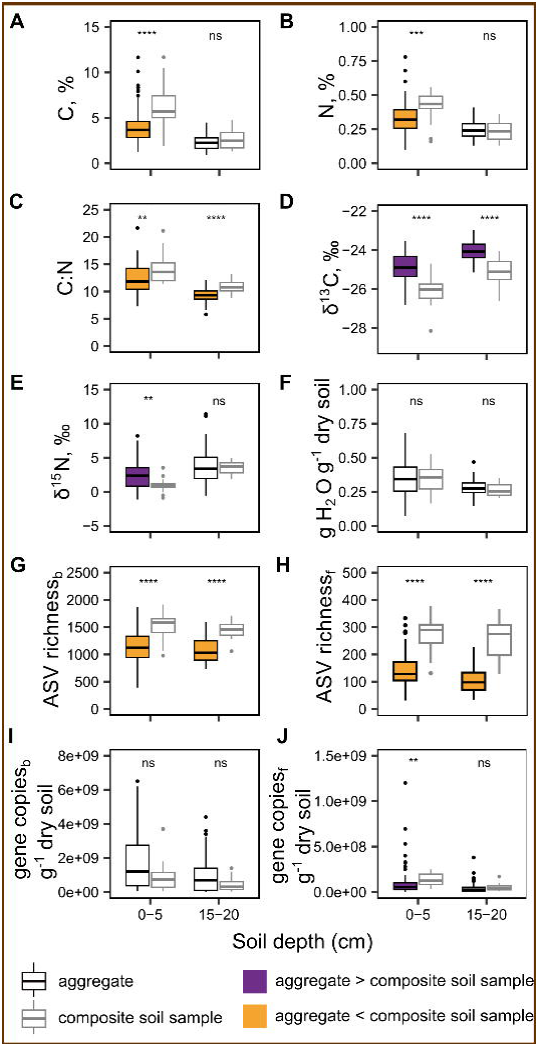
Abiotic and microbial variables partly differed between aggregates and composite soil samples. Boxplots display (**A**) carbon content (C, %) and (**B**) nitrogen content (N, %), (**C)** carbon-to-nitrogen ratio (C:N), isotopic ratios of (**D**) natural carbon (δ^13^C, ‰) and (**E**) nitrogen (δ^15^N, ‰), (**F**) soil water content (g H_2_O g^−1^ dry soil), (**G**) bacterial and archaeal ASV richness (ASV richness_b_), (**H**) fungal ASV richness (ASV richness_f_), (**I**) bacterial and archaeal gene copies (16S rRNA gene copies g^−1^ dry soil), and (**J**) fungal gene copies (ITS gene copies g^−1^ dry soil) for aggregates and composite soil samples in 0-5 and 15-20 centimetres soil depth. Black-border boxes represent aggregates, grey-bordered boxes represent composite soil samples, and points depict outliers. The number of measurements per variable boxplots were based on varies and depends on the specific variable (0-5 cm: between 64 and 92 aggregates, 14-20 composite soil samples; 15-20 cm: 73-92 aggregates, 18-20 composite soil samples; see exact number of measurements underlying boxplots in Table S3). The fill colour of aggregate boxes indicates whether the variable was significantly larger or smaller in aggregates compared to composite soil samples. Asterisks above the boxplots indicate the significance of differences between aggregates and composite soil samples for each soil depth, calculated using a Kruskal Wallis test (**** P value≤0.0001, *** P value≤0.001, ** P value ≤0.01, * P value ≤0.05, ns P value>0.05).

Carbon and nitrogen content, stoichiometric and isotopic ratios, and soil water content of aggregates and composite soil samples differed between the two soil depths (Fig. 2A-F). Samples from 15-20 centimetres soil depth were characterised by lower carbon and nitrogen content, narrower carbon-to-nitrogen ratio, more positive ^13^C and ^15^N value, and slightly lower soil water content compared to samples from 0-5 centimetres soil depth.

#### 3.1.3 Fungal density was higher in composite soil than in individual aggregates

We calculated 16S rRNA genes and ITS gene copy numbers using ddPCR to compare microbial density between 2-millimetre-sized aggregates and composite soil samples. We observed significantly higher fungal gene copy numbers per gram dry soil in composite soil samples than in aggregates (Fig. 2J, 0-5 cm: χ^2^=7.6, P value=0.006, 15-20 cm: χ^2^=3.5, P value=0.06). In contrast, bacterial and archaeal gene copy numbers per gram dry soil did not differ significantly between the two sample types. However, they were slightly higher in aggregates than composite soil samples (Fig. 2I, 0-5 cm: χ^2^=3.1, P value=0.08, 15-20 cm: χ^2^=1.7, P value=0.2).

#### 3.1.4 Bacterial, archaeal, and fungal richness was higher in composite samples than individual aggregates

We determined number of unique microbial ASVs of individual aggregates and composite soil samples after we rarefied samples to the same number of reads for bacteria and archaea and fungi, respectively, to guarantee equal sequencing depth. We detected significantly less bacterial, archaeal, and fungal ASVs in 2-millimetre-sized aggregates than in composite soil samples (Fig. 2G, 0-5 cm: χ^2^=2.4, P value≤0.0001, 15-20 cm: χ^2^=36.3, P value≤0.0001; Fig. 2H, 0-5 cm: χ^2^=34.9, P value≤0.0001, 15-20 cm: χ^2^=35.2, P value≤0.0001).

### 3.2 Relationships among microbial and organic matter properties were scale-dependent

#### 3.2.1 Organic matter properties were more strongly correlated in composite soil samples than in aggregates

We calculated Spearman rank correlations among and between abiotic variables and microbial variables of the two sample types, respectively, to test whether and how (strongly) variables are related in aggregates and in composite soil samples. Our data shows that most abiotic and microbial variables were correlated in the same way across spatial scales, however relationships differed in their strength between the two sample volumes (Fig. 3A-3F). We found carbon and nitrogen content, carbon-to-nitrogen ratio, and δ^13^C and δ^15^N to be negatively correlated in both aggregates and composite soils (Fig. 3A, 3B, 3D, 3E). Yet, organic matter properties were more strongly correlated (i.e. had a more positive or more negative correlation coefficient) in composite soil samples than in aggregates (Fig. 3C, 3F). In contrast, soil water content was correlated more strongly with organic matter properties and microbial community properties in aggregates compared to composite soil samples in topsoil (Fig. 3C).

**Fig. 3.**
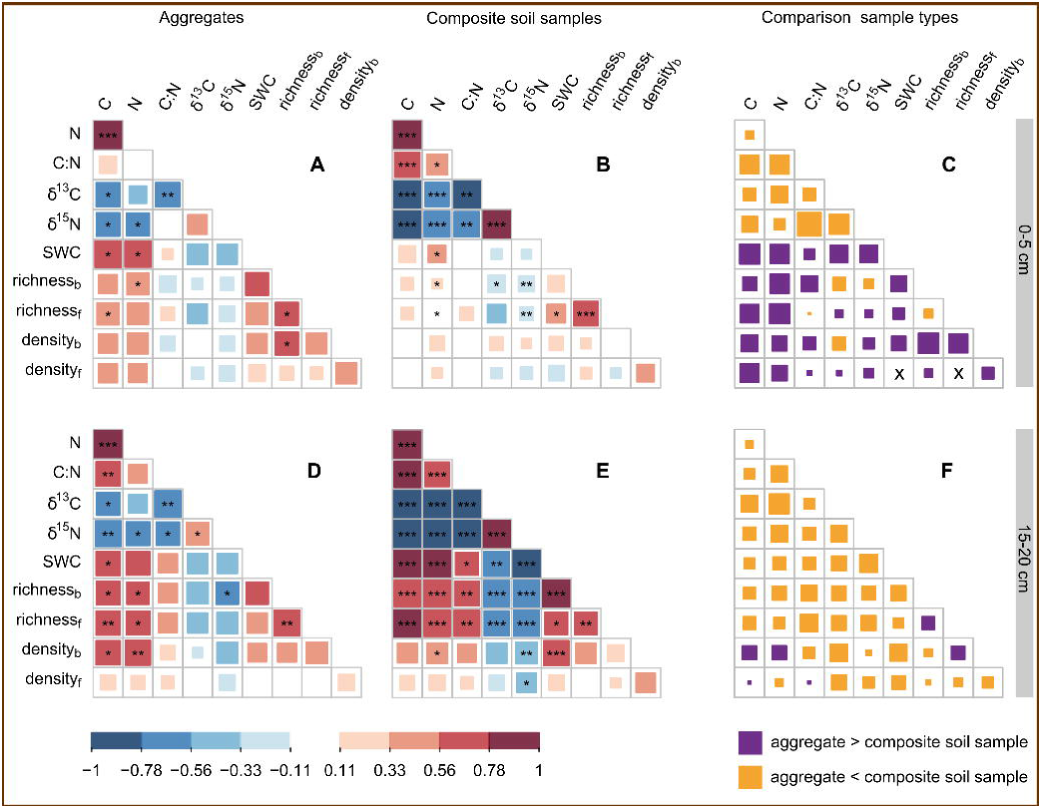
Correlations among organic matter properties, soil moisture, and microbial community parameters differed between aggregates and composite soil samples. (**A, D**) Correlation heatmaps show the median pairwise Spearman rank correlation coefficients for carbon (C, %), nitrogen content (N, %), carbon-to-nitrogen ratio (C:N), δ^13^C (‰), δ^15^N (‰), soil water content (SWC, g H_2_O per g dry soil), bacterial and archaeal ASV richness (richness_b_), fungal ASV richness (richness_f_), 16S rRNA gene copies (density_b_, gene copy numbers per g dry soil), and ITS gene copies (density_f_, gene copy numbers per g dry soil) in 2-mm-sized aggregates from 0-5 and 15-20 centimetres soil depth. Median correlation coefficients were calculated from individual correlation coefficients of 5000 aggregate subsets, each containing 20 randomly picked aggregates (one aggregate per soil core). Asterisks depict the median P values for pairwise correlations of all 5000 aggregate subsets (*** P value≤0.001, ** P value≤0.01, * P value≤0.05). Median P values are not directly associated with the median correlation coefficient displayed in the graphs. (**B, E**) Correlation heatmaps depict the pairwise Spearman rank correlation coefficients for composite soil samples, with asterisks indicating the significance of correlations (*** P value≤0.001, ** P value≤0.01, * P value≤0.05). (**A, B, D, E**) Colour indicates the direction of correlations (positive or negative), while the shade of the colour and the size of the squares indicate correlation strength. (**C, F**) Plots display the absolute differences in Spearman correlation coefficients between aggregates and composite soil samples, which were determined by subtracting the correlation coefficient of composite soil samples from the median correlation coefficient of aggregates. Differences were only calculated between correlation coefficients with the same sign. The colour of the squares indicates whether correlations were stronger in aggregates than in composite soil samples or vice versa; “x” marks indicate relationships where the sign of the correlation differed between aggregates and composite soil samples.

#### 3.2.2 Microbial and organic matter properties were more strongly correlated in aggregates than in composite soil samples in the topsoil

Comparing Spearman rank correlation coefficients between aggregates and composite soil samples, we found that in topsoil, microbial richness and bacterial and archaeal density were correlated with carbon and nitrogen content considerably stronger within aggregates than within composite soil samples (Fig. 3C). In 15-20 centimetres soil depth, however, microbial richness was stronger correlated with abiotic parameters in composite soil samples compared to aggregates, while bacterial density was stronger correlated with soil carbon and nitrogen content in aggregates (Fig. 3F). Fungal density, overall, was only weakly and non-significantly correlated with the other investigated variables in both sample volumes.

Including all available aggregates (n=92 for 0-5 cm soil depth, n=98 for 15-20 cm) in the correlation analysis (Fig S1) resulted in similar correlation coefficients for each pairwise correlation as the median correlation coefficients of 5000 combinations of 20 randomly collected aggregates (Fig. 3A, 3D). Differences in correlation strengths (i.e. correlation coefficients) between aggregate and composite soil samples were thus the same in the full-aggregate dataset compared to the analysis based on the median of 5000 20-aggregate subsets (Fig. 3C, 3F, S2C, S2F). However, all P value of correlations were far more significant (i.e. had lower P values) when determined for 92 or 98 aggregates compared to the median P values of 5000 P value 20-aggregate subsets (Fig S1).

## 4 Discussion

Soils are highly heterogeneous habitats that exhibit large spatial variability in microbial distribution and environmental conditions down to the micrometre-scale (Young et al., 2008; Vos et al., 2013; Smercina et al., 2021). According to scale theory, it can be expected that observation scale affects research findings in soil and that relationships among variables are scale-dependent. Consequently, observed relationships would not be transferrable between samples that differ in sample grain, in other words, sample size. In this study, we aimed to test whether relationships among microbial community, organic matter properties, and soil water content differ between individual 2-millimetre-sized soil aggregates and composite soil samples, two sample types that differ pronouncedly in soil volume (Fig. S1). Additionally, we assessed whether 2-millimetre-sized soil aggregates represent a special component of soil and offer particular environmental conditions that differ from other areas of the soil matrix. Our results show that individual 2-millimetre-sized aggregates and composite soil samples differed systematically in microbial community and organic matter properties. Aggregates were systematically enriched in ^13^C and exhibited lower carbon-to-nitrogen ratio than composite soil samples in both soil depths. Additionally, they contained proportionally less carbon and nitrogen than composite soil samples in topsoil. Moreover, we found that relationships among and across microbial and organic matter parameters depended on sample volume.

Carbon-to-nitrogen ratio and δ^13^C can be used as proxies for the recycling status of organic matter in soil (Zechmeister-Boltenstern et al., 2015; Lorenz et al., 2020). This is because microbial respiration discriminates against the heavier carbon isotope (^13^C), which leads to microbial biomass being enriched in ^13^C compared to their food. At the same time, carbon-to-nitrogen ratio is more narrow in microbial biomass than in plant tissue (Mooshammer et al., 2014; Zechmeister-Boltenstern et al., 2015). Consequently, soil organic matter has a higher δ^13^C value and a lower carbon-to-nitrogen ratio the higher its ratio of microbial-derived to plant-derived organic matter is (Dijkstra et al., 2006). Additionally, δ^13^C values of organic matter increase the more often carbon has been recycled by microbes. Gradients in carbon-to-nitrogen ratio and ^13^C-enrichment thus indicate a shift from plant-derived organic matter to more decomposed and microbial-derived, and eventually, also more-often recycled organic matter (Rumpel et al., 2002; Rumpel and Kögel-Knabner, 2011). The higher δ^13^C and lower carbon-to-nitrogen ratio that were observed in aggregates suggest that proportionally less fresh or sparsely decomposed plant litter was present in 2-millimetre-sized soil aggregates than in other soil components. The lower organic matter content (carbon and nitrogen content) of aggregates from the top five centimetres compared to composite soil samples from the same layer indicates that they contained relatively smaller fractions of carbon- and nitrogen-dense material (such as aboveground plant residuals) compared to other areas of the soil matrix. Bioturbation, soil mixing via the activity of soil fauna, strongly impacts the amount of aboveground plant residual input to soil (Tonneijck and Jongmans, 2008). Burrows created by large soil fauna like earthworms might pose areas of fracture along which soil will break upon soil sampling and subsequent soil disintegration. Consequently, the 2-millimetre-sized aggregates which we would sample would not contain fresh burrows and associated plant material. The diminished difference in organic matter content between aggregates and composite soil samples in the deeper soil layer might be the result of the general decline in aboveground plant residuals with soil depth due to the increasing distance to the soil surface and reduced bioturbation with soil depth (Tonneijck and Jongmans, 2008; Wilkinson et al., 2009).

Lower organic matter content in aggregates compared to composite soil samples in the topsoil layer could also indicate lower occurrence of fungal hyphae and/or plant roots in aggregates than other components of the soil matrix. Overall, fungal hyphae and roots are substantial carbon sources in soil (Prescott and Vesterdal, 2021). In fact, aggregates were less densely populated by fungi than other regions of the soil matrix. At the same time, we detected slightly higher bacterial and archaeal density in aggregates than in composite soil samples. These opposing trends in fungal and bacterial/archaeal biomass suggest that 2-millimetre-sized soil aggregates offered physico-chemical conditions that were distinct from other components of the soil matrix. Differences in the extent of fungal colonisation might indicate different pore size distribution in aggregates and other components of the soil matrix (smaller pores within and larger pores between aggregates), a view that has been challenged recently (Vogel et al., 2022). However, our results are consistent with studies that showed that fungal hyphae more readily colonise larger pores (Otten et al., 2004) and rarely occur in micropores (Killham, 1994), whereas bacteria are proposed to be more abundant in smaller pores where they are better sheltered from microbial predators (soil fauna) (Maier et al., 2009; Wright et al., 2014).

Despite the differences in individual variables between soil aggregates and composite soil samples, both sample types depicted lower variability of chemical composition in the lower soil layer than the topsoil, indicating decreased spatial heterogeneity in chemical properties with soil depth. This reduced spatial heterogeneity might have caused the observed lower variability of microbial density across aggregates and composite soil samples, respectively, in the deeper soil layer compared to the topsoil.

Our analysis showed that relationships among and between abiotic and biotic variables depended on soil volume. Strength of correlations differed between individual 2-millimetre-sized aggregates and composite soil samples. Organic matter properties were more strongly correlated in composite soil samples than in aggregates. This finding matched our expectation, namely that correlation strength increases with increasing soil volume. Producing composite soil samples averages out the small-scale heterogeneity of the soil matrix, reduces noise and by that potentially strengthens correlations (Wiens, 1989; Wu and Li, 2006).The stronger correlations among organic matter properties which were observed for composite soil samples compared to aggregates might be the mere result of differences in sample volume.

Correlations among microbial community variables and organic matter properties in topsoil showed an opposing trend. Variables correlated weaker in composite soil samples than in aggregates. We propose two possible explanations for the observed pattern. (i) Variables scaled differently with soil volume. While carbon content might scale linearly with soil volume, microbial taxon (ASV) richness might not. Consequently, the link observed in small-sized soil units become weaker as soil volume increases. (ii) A second possible explanation is that soil is composed of a variety of components which differ in environmental conditions. 2-millimetre-sized soil aggregates seem to be stable for long enough timeframes to allow the establishment of relationships between microbes and their immediate environment. In contrast, other areas of the soil matrix might be temporally less stable and hence exhibit weaker relationships between microbes and environmental parameters than aggregates. Soils may include areas which are exposed to disturbance or mixing more often than aggregates, which hampers or destroys a link between microbes and environmental properties. Some areas of the soil matrix may contain fresh particulate organic matter but exhibit low microbial density as microbes might not yet have started growing in response to newly available substrate. We believe that the weaker correlations among microbial and organic matter properties in the topsoil are more likely the result of the soil matrix being composed of various areas/components which differ in their characteristics than being the mere result of sample volume. After all, aggregates differed pronouncedly from composite soil samples in organic matter properties in topsoil.

In contrast, in the deeper soil layer microbial communities and organic matter properties were similarly strong correlated in aggregates and composite soil samples except for the correlation between bacterial density and organic matter content. Bacterial density was more strongly correlated with carbon and nitrogen content in aggregates than composite soil samples in both soil layers. This underpins that there are other soil areas than aggregates where microbes are less closely related with organic matter availability than in aggregates also for the deeper soil layer. Notably, correlations were overall stronger in the deeper soil layer than in the topsoil. This might be the result of environmental properties becoming more homogenous with increasing soil depth, which could strengthen correlations due to decreased variability in parameters.

We observed a large variability in pairwise Spearman correlation coefficients and P values across the 5000 aggregate subsets that included an arbitrary collection of 20 aggregates for both soil depths. Whereas medians of correlation coefficients of aggregate subsets (Fig. 3) approximated correlation coefficients that were calculated for the full dataset of 90 to 100 aggregates (Fig. S2), P values were notably more significant in the case of 90 or 100 aggregates than averages or medians of 5000 20-aggregates subsets. In addition, the high underlying variability of correlation coefficients across the 5000 20-aggregate subsets demonstrates that correlations obtained from any one 20-aggregate dataset would have a large random component and may not represent underlying patterns correctly. Hence, we conclude that when decreasing sample volume down to the aggregate size, it is highly recommendable to increase sample numbers accordingly.

Our results indicate that composite soil samples were not blends of only 2-millimetre-sized soil aggregates but contained something(s) else besides 2-millimetre-sized soil aggregates. The soil matrix seemed to be composed of various components which vary in chemical properties. Inversely, this means that 2-millimetre-sized soil aggregates cannot be interpreted as small-volume composite soil samples but should be considered as special components of the soil matrix.

## 5 Conclusion

In this study, we have demonstrated that the choice of observation scale can substantially affect research outcomes in soil. Our study underpins that it is crucial to identify the appropriate sampling scale for a specific research question. Based on our findings and theoretical considerations, we recommend that one needs to study small-volume sample units to elucidate relationships between microbial communities and their environment. If we had solely examined composite soil samples, we would have underestimated or even missed the link between organic matter properties and microbial community parameters in topsoil. We have shown that soil aggregates are a suitable sampling unit to study these relationships. Individual millimetre-sized soil aggregates have allowed insights into relationships among microbial community parameters and environmental properties. However due to the large observed variability across aggregate subsets which contain only a small number of aggregates, it is advisable to investigate a relatively large number of aggregates. Moreover, our data indicates that soil is composed of a variety of components which vary in environmental conditions. Our findings indicate that 2-millimetre-sized soil aggregates are special components of soil and systematically differ in habitat characteristics like recycling status from other areas of the soil matrix. We propose that individual soil aggregates carry additional information of composite soil samples and will improve our understanding about soil as a microbial habitat.

## 7 Acknowledgments

This research was funded by the European Research Council under the European Union’s Horizon 2020 research and innovation programme (grant agreement No 819446). We thank Barbara Kitzler for enabling access to the field site, Julia Mor, Kian Jenab, and Christian Ranits for helping with processing and measuring samples in the lab, Margarete Watzka for measuring organic matter properties of samples, Petra Pjevac for advising and coordinating DNA extraction and amplicon sequencing of samples, Bela Hausmann for downstream amplicon sequencing data processing, and Joana Seneca for submission of amplicon sequencing data to NCBI.

## 8 CRediT authorship contribution statement

**Eva Simon**: Formal analysis, Investigation, Visualization, Methodology, Writing – original draft, Writing – review & editing. **Ksenia Guseva**: Investigation, Writing – review & editing. **Lauren Alteio**: Investigation, Writing – review & editing. **Christina Kaiser**: Investigation, Conceptualization, Funding acquisition, Supervision, Writing – review & editing.

## 9 Data availability

Raw 16S rRNA genes and ITS2 amplicon sequencing data have been deposited in the NCBI Sequence Read Archive under the BioProject accession number PRJNA1090291. Soil organic matter properties and soil water content are available on the Zenodo repository (https://zenodo.org/records/14929829).

