## Supplementary Material for "Strengths of relationships among soil microbial and organic matter properties are scale-dependent"

**Table S1 PCR cycling conditions used for amplicon sequencing of 16S rRNA genes and ITS2 regions.** PCR reactions (25 µl) comprised 1x DreamTaq Green PCR master mix, 0.1 µg µl-1 BSA, 0.25 (for 16S rRNA) or 0.5 (for ITS2) µmol l-1 of each primer and 2 µl of DNA template.

|  | **Temperature (°C)** | **Time** |
| --- | --- | --- |
| **ITS2,** ISOF-T/ITS4 | 94 | 4 min |
|  | 94 | 45 sec (35 cycles) |
|  | 48 | 45 sec |
|  | 72 | 1 min |
|  | 72 | 10 min |
| **ITS2,** Nested PCR (gITS7/ITS4) | 94 | 4 min |
|  | 94 | 30 sec (20 cycles) |
|  | 53 | 30 sec |
|  | 72 | 30 min |
|  | 72 | 10 min |
| **16S** V4 (515F/806R) | 94 | 3 min |
|  | 94 | 45 sec (30 cycles) |
|  | 52 | 60 sec |
|  | 72 | 60 min |
|  | 72 | 10 min |

**Table S2 Primers and PCR cycling conditions for the thermocycler used for Digital Droplet PCR to amplify 16S rRNA genes and ITS1 regions within droplets.** PCR reactions (22 µl) consisted of 11 µl QX200 ddPCR EvaGreen Supermix (1x), 0.2 µl of forward primer (0.1 µM) and 0.2 µl of reverse primer (0.1 µM), 8.6 µl DNase-free water and 2 µL DNA template. Adjustments were made to the ratio between DNA template and reagents when additional DNA template was required to achieve an optimal DNA concentration of 0.1 (for 16S rRNA) and 0.25 (for ITS1) ng/µl for PCR reactions.

| **16S rRNA genes** (515F (Parada et al., 2016) and 806R (Apprill et al., 2015)) | **ITS1 regions** (ITS1F & ITS2 (White, T. J. et al., 1990)) |
| --- | --- |
| 95°C for 5 min | 95°C for 5 min |
| 5 cycles of 95 °C for 30 s and 57°C for 2.5 min (−1 °C each step) | 5 cycles of 95°C for 30 s and 60°C for 2.5 min (−1°C each step) |
| 35 cycles of 95°C for 30 s and 52°C for 2.5 min | 35 cycles of 95°C for 30 s and 55°C for 2.5 min |
| 4°C for 5 min | 4°C for 5 min |
| 90°C for 5 min | 90°C for 5 min |
| 10°C hold | 10°C hold |
| Storage at 4°C (min 1 h) before droplet reading | Storage at 4°C (min 1 h) before droplet reading |

**Table S3 Number of measurements per analysed variable for aggregates and composite soil samples of both soil depths (0-5, 15-20 centimetres soil depth).** C – carbon, N – nitrogen, SWC – soil water content

| **Variables** | **Aggregates** | | **Composite soil samples** | |
| --- | --- | --- | --- | --- |
|  | **0-5 cm** | **15-20 cm** | **0-5 cm** | **15-20 cm** |
| C content | 92 | 98 | 20 | 20 |
| N content | 92 | 98 | 20 | 20 |
| C-to-N ratio | 92 | 98 | 20 | 20 |
| δ^13^C | 92 | 98 | 20 | 20 |
| δ^15^N | 92 | 98 | 20 | 20 |
| SWC | 64 | 73 | 20 | 20 |
| 16S ASV richness | 90 | 96 | 19 | 20 |
| ITS ASV richness | 87 | 96 | 20 | 18 |
| 16S gene copies | 75 | 86 | 19 | 19 |
| ITS gene copies | 83 | 77 | 14 | 18 |

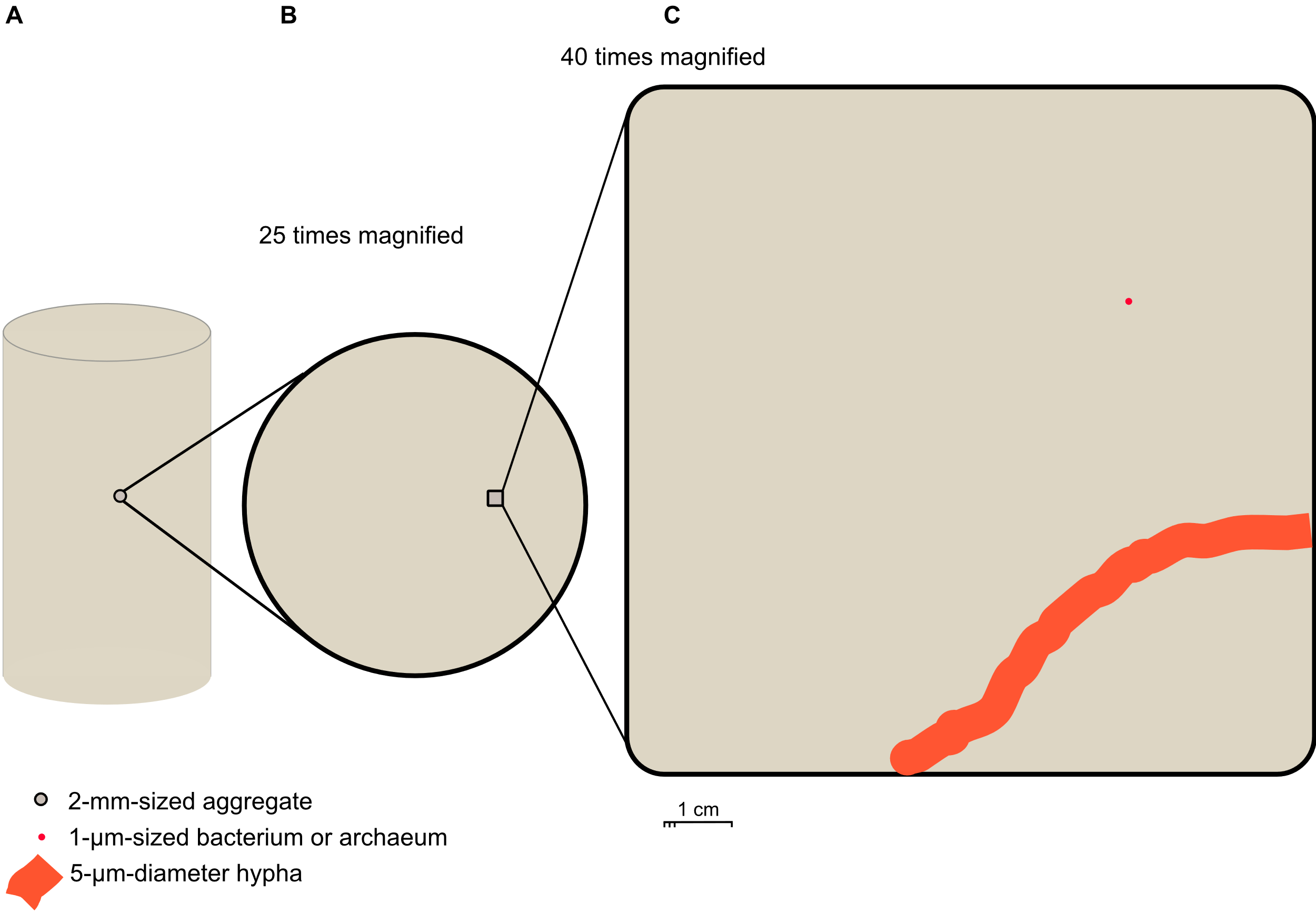

**Fig. S1 Schematic representation of the size differences between a soil core, a 2-millimetre-sized soil aggregate, a bacterial/archaeal cell, and a fungal hypha.** (**A**) shows a 2-millimetre-sized soil aggregate in a soil core (diameter=3 cm, height=5 cm) that comprises several cubic centimetres of soil. When sieved, the soil core produces a composite soil sample. The figure illustrates the significant difference in the soil volume represented by the two sample types, soil core/composite soil sample and individual soil aggregate. (**B**) depicts a 2-mm-sized soil aggregate which is magnified 25 times. (**C**) pictures a section of the 2-mm-sized aggregate, further magnified 40 times, alongside a 1 micrometre (µm)-sized bacterium or archaeum, and a fungal hypha of 5 µm diameter. The aggregate section, bacterial/archaeal cell, and fungal hypha are all magnified 1000 times. It has been reported that over 60% of bacterial and archaeal cells in various soils are smaller than 1.2 µm (Portillo et al., 2013), whereas the diameter of fungal hyphae ranges from 1 and 30 µm (Islam et al., 2017).

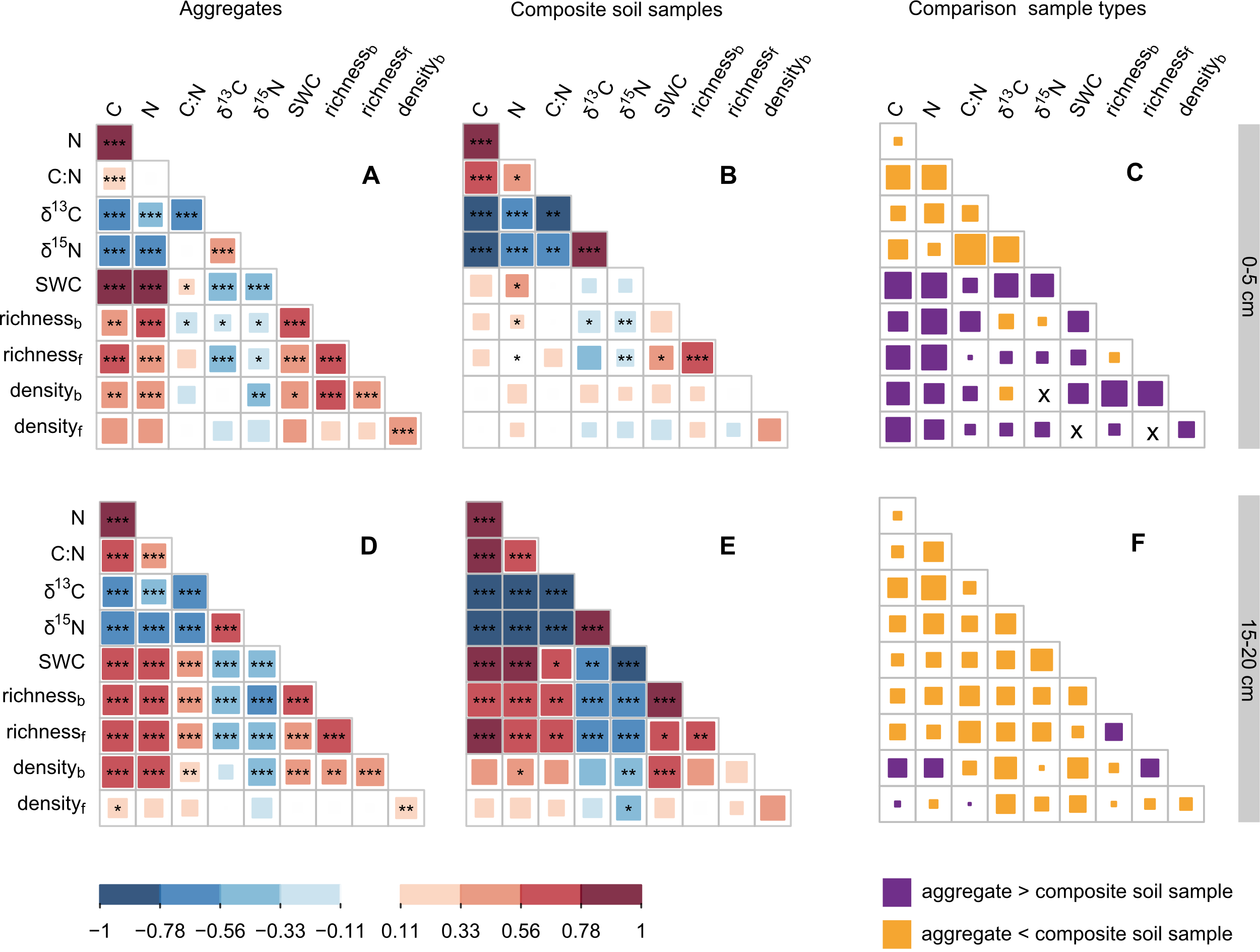
**Figure S2 Correlation heatmaps show correlations among organic matter properties, soil moisture, and microbial community parameters for aggregates (n=190) and composite soil samples (n=40).** Correlation heatmaps depict the pairwise Spearman rank correlation coefficients of carbon (C, %), nitrogen content (N, %), carbon-to-nitrogen ratio (C:N), δ^13^C (‰), δ^15^N (‰), soil water content (SWC, g H_2_O per g dry soil), bacterial and archaeal ASV richness (richness_b_) and fungal ASV richness (richness_f_), 16S rRNA gene copies (density_b_, gene copy numbers per g dry soil) and ITS gene copies (density_f_, gene copy numbers per g dry soil) for (**A, D**) 2-mm-sized aggregates and (**B, E**) composite soil samples (see number of measurements for each variable in Table S3). (**A, B, D, E**) Asterisks give significance of correlations (*** P value≤0.001, ** P value ≤0.01, * P value ≤0.05). The colour of the squares indicates direction of correlations (positive or negative), while the shade of the colour and the size of the squares indicate correlation strength. (**C, F**) Plots display the absolute differences in Spearman correlation coefficients between aggregates and composite soil samples, which were determined by subtracting the correlation coefficient of composite soil samples from the correlation coefficient of aggregates. Differences were only calculated between correlation coefficients with the same sign. The colour of the squares indicates whether correlations were stronger in aggregates than in composite soil samples or vice versa; “x” marks indicate relationships where the sign of the correlation differed between aggregates and composite soil samples.
